# Highly reliable, targeted photothermal cancer therapy combined with thermal dosimetry using indocyanine green lactosome

**DOI:** 10.1101/659334

**Authors:** Shinsuke Nomura, Yuji Morimoto, Hironori Tsujimoto, Manabu Harada, Daizoh Saitoh, Isao Hara, Eiichi Ozeki, Ayano Satoh, Eiji Takayama, Kazuo Hase, Yoji Kishi, Hideki Ueno

**Author notes:** Corresponding author Yuji Morimoto, Department of Physiology, National Defense Medical College, 3-2, Namiki, Tokorozawa, Saitama 359-8513, Japan.

## Abstract

Indocyanine green (ICG) is a near-infrared light-absorbing substance. Thus, when a tumor in which ICG has accumulated is irradiated with a near-infrared (NIR) laser, only the tumor can be heated by a photothermal reaction. We developed ICG lactosome, a novel drug delivery system (DDS) composed of polymeric micelles and ICG that shows selective accumulation in tumor based on an enhanced permeability and retention (EPR) effect. We showed that ICG lactosome accumulated in a tumor by using an intradermal tumor mouse model of a murine colon cancer cell line (Colon26) transfected with Nano lantern luminescent protein (NLC26). Two days after the administration of ICG lactosome, the tumor was irradiated with an 808-nm diode-laser while monitoring tumor temperature. The results showed that the treated tumors were cured when the peak of tumor temperature during NIR irradiation reached 43°C or higher. To verify these results, photothermal therapy (PTT) using ICG lactosome was carried out using a newly developed system that can control the temperature at the NIR irradiation site at a constant level. All of the tumors that had been kept at 43°C during irradiation were cured, while 2 of 5 tumors that had been kept at 42°C were not cured, and none of tumors that had been kept at a temperature below 41°C were cured. ICG lactosome-assisted PTT combined with thermal dosimetry is a highly reliable method for cancer treatment and may afford further clinical opportunities for PTT.

## Introduction

Indocyanine green (ICG), which was approved by the U.S. Federal Drug Administration (FDA) in 1959 (Zheng et al., 2014), has been widely used clinically not only for evaluation of liver function and cardiac output but also as a fluorescent biomedical imaging tool for identification of sentinel lymph nodes of breast cancer (Toh et al., 2015) and gastric cancer (Okubo et al., 2018) and for real-time evaluation of blood flow in organs (Li et al., 2018, Milstein et al., 2016). ICG is also known as a near-infrared (NIR) light-absorbing substance, that is a photo thermal agent, and thus when a tissue in which ICG exists is irradiated with NIR light, only the tissue irradiated can be heated by a photo thermal reaction (Liu et al., 2002). Studies on anti-cancer treatment using this phenomenon have also been performed (Chen et al., 1995a, Chen et al., 1995b), and a growth inhibitory effect was found in tumor tissue in which ICG was locally injected and then the tissue was irradiated with NIR light (Chen et al., 1996). However, the properties of ICG have some limitations such as poor photostability, non-specific targeting, and short half-life. In order to overcome these problems and to deliver more ICG to the tumor, many researchers have attempted to develop an ICG-based drug-delivery system (DDS) to exert an enhanced permeabilit and retention (EPR) effect (Matsumura and Maeda, 1986, Wang et al., 2018). Various ICG nanoparticles including lipid-based, polymer-based, and mesoporous silica ICG nanoparticles have been developed (Wang et al., 2018). Subcutaneous tumors have been successfully eradicated by a photothermal reaction using ICG nanoparticles (Chen et al., 2016, Zheng et al., 2014).

In our previous study, we established ICG lactosome from self-assembly of poly (sarcosine)–poly (L-lactic acid) (PS–PLLA) biodegradable amphiphilic block copolymers loaded with an ICG derivative (Makino et al., 2009). Since ICG lactosome particles are about 30-40 nm in diameter, ICG lactosome shows preferable accumulation in the tumor due to the EPR effect (Tsujimoto et al., 2015b, Tsujimoto et al., 2014b).When tumors in which ICG lactosome had accumulated were exposed to NIR light, the proliferation of tumor cells was greatly inhibited mainly due to the photothermal reaction: the therapeutic effectiveness of ICG lactosome has been shown in peritoneal carcinomatosis (Tsujimoto et al., 2014b) and metastatic lymph nodes (Tsujimoto et al., 2015b) of gastric cancer, spinal metastasis of breast cancer (Funayama et al., 2012, Funayama et al., 2013, Tsukanishi et al., 2014), and hepatocellular carcinoma (Tsuda et al., 2017). However, most of the irradiation parameters such as irradiation time and fluence rate were predetermined before NIR irradiation in those studies, and the optimal therapeutic effects have not been determined. Furthermore, the photothermal efficacy of ICG lactosome as a NIR light-absorbing agent has not been demonstrated.

We have conducted animal experiments on phototherapy using ICG lactosome, but the expected therapeutic effect was not obtained for all. Therefore, in order to obtain a reliable therapeutic effect, we examined the relationship between photothermal therapy (PTT) parameters and therapeutic effects. We found that the maximum temperature of the tumor during NIR irradiation was most strongly correlated with the therapeutic effect.

## RESULTS

### Outline of the experimental design

An intradermal tumor model was used to verify the anti-tumor effect of PTT using ICG lactosome on a tumor. In the tumor model, the only tissue between the tumor and the laser was epidermal tissue. Based on the results of our previous experiments, ICG lactosome was administered 48 h before NIR laser irradiation. Fluorescence images were obtained by an *in vivo* imaging system (IVIS; PerkinElmer) before NIR laser irradiation. The temperature of the tumor during NIR irradiation was monitored in real time by using an infrared radiation thermometer (FT-H10; Keyence). On day 21, the three-dimensional size of the tumor was measured by using a digital caliper.

### Specific accumulation of ICG lactosome in the tumor

In order to determine whether ICG lactosome accumulates specifically in tumor tissue, the consistency between the fluorescence and the actual tumor location from ICG lactosome was confirmed using a Colon26 cell line transfected with Nano-lantern luminescent protein (NLC26), which degrades the luminescent substrate coelenterazine h (Wako Pure Chemical Industries Ltd., Osaka, Japan). Figure 1A shows the consistency among the white image (WLI), the bioluminescent image emitted from a luminescent substrate degraded in tumor cells (BLI), and the fluorescent image from ICG lactosome (FLI). These images were taken at 48 hours after injection by the IVIS. Fluorescence images were taken at 5 min, 30 min, 4 h, 12 h, 24 h and 48 h after intravenous injection of ICG lactosome for observing tumor selectivity (Fig. 1B). At 12 h after ICG lactosome injection, the margin of the tumor started to become clear in the fluorescent image. After that, the tumor/non-tumor tissue fluorescence intensity ratio increased, and the ratio was more than 12-times higher at 48 h.

**Fig. 1.**
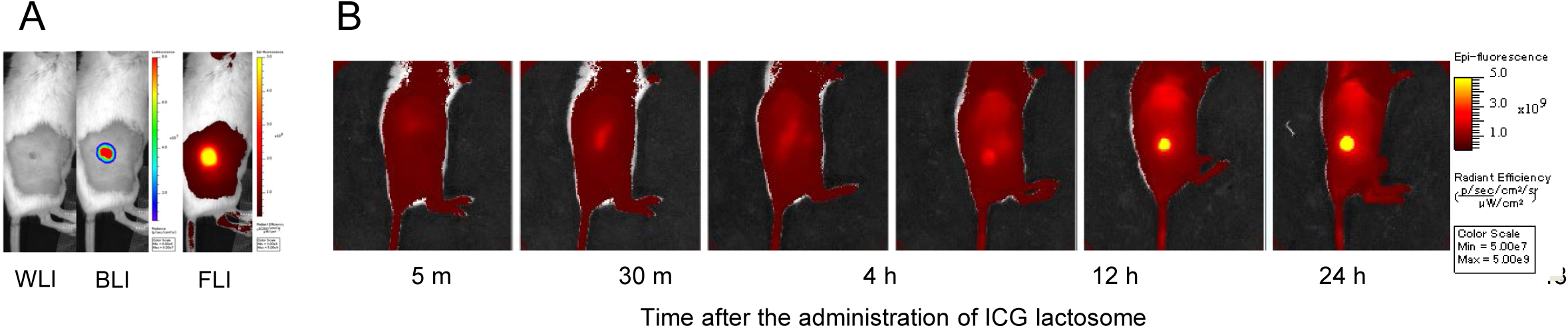
*In vivo* imaging before and after irradiation. Representative examples of a white light image (WLI), bioluminescence image (BLI), and ICG lactosome fluorescence image (FLI) before PTT. B. Time-lapse image of fluorescence from ICG lactosome. The location of fluorescence being consistent with the tumor location was identified 12 h after the injection of ICG lactosome, and fluorescence at the tumor gradually intensified with time: the contrast ratio (= fluorescence at tumor site /fluorescence at non-tumor site) reached more than twelve at 48h after administration.

Since fluorescence from ICG lactosome consistent with luminescence from the tumor was observed, ICG lactosome had excellent selective accumulation in the tumor.

### Correlation of temperature in PTT with antitumor effect

Intradermal tumor model mice were used to investigate changes in the tumor temperature and anti-tumor effect with changes in the parameters of NIR irradiation. First, we changed the fluence rate to observe the time courses of tumor temperature and antitumor effect. Fluence rates were set at 250, 500, 750 and 1000 mW/cm^2^. The irradiation time was set at 1000 s and the dose of ICG lactosome was 8.8 mg/kg. When the NIR laser was turned on, the temperature at the surface of the skin covering the tumor reached a peak at about 120 s at all fluence rates (Fig. 2A). After reaching the peak, the temperature remained constant or gradually decreased. When the fluence rate was increased from 250 to 1000 mW/cm^2^, the temperature during NIR irradiation increased (Fig. 2A); however, there was no statistically significant difference in tumor temperature between NIR irradiation with 750 mW/cm^2^ and that with 1000 mW/cm^2^. After NIR irradiation, we investigated how the fluence rate changes antitumor effect (Fig. 2B). Significant differences were observed between a fluence rate of 1000 mW/cm^2^ and fluence rates of 0, 250 mW/cm^2^, and between a fluence rate of 750 mW/cm^2^ and 0 mW/cm^2^. Although tumor growth seemed to be suppressed in proportion to the fluence rate, there were no significant differences among the three fluence rates (250, 500 and 750 mW/cm^2^).

**Fig. 2.**
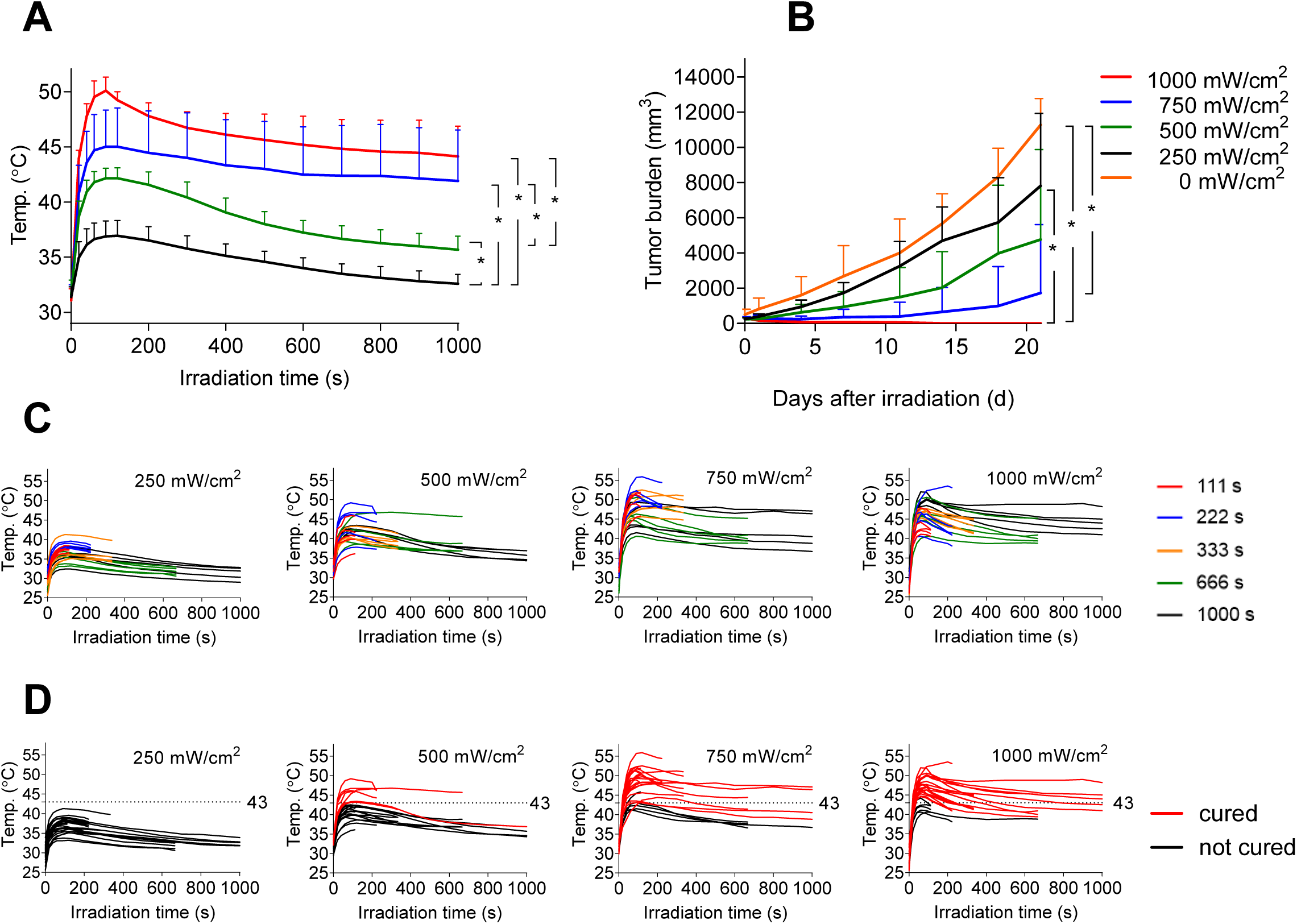
A. Changes in temperature during NIR irradiation of the tumor at each fluence rate. (*P < 0.05, n = 5 each) B. Corresponding growth curves of tumors in mice treated with PTT at each fluence rate for 1000 s (*P < 0.05, n = 5 each) C. Raw profile of temperature change in each individual tumor during NIR irradiation. D. The temperature curves in “C” are color-coded into two groups. The tumors for which the temperature curves are shown in red were eradicated (cured), and the tumors for which the temperature are shown in black were not cured (proliferation of tumor cells).

Next, to verify the relationship between anti-tumor effect and time of NIR irradiation, we examined the anti-tumor effect at irradiation times of 111, 222, 333, 666 and 1000 s with a constant fluence rate (either at 250, 500, 750 or 1000 mW/cm^2^) (Fig. 2C). At any fluence rate, there was no correlation between tumor size on day 21 and irradiation time (Fig. S1). When analyzing the same data in another way, tumor size at each irradiation time did not correlate with fluence rate (Fig. S2). Analysis of the time course of temperature at each fluence rate revealed that the tumor disappeared when the temperature of the tumor increased above 43°C (Fig. 2D, 3): all but two of the 75 tumors disappeared regardless of fluence rate and irradiation time. Figure 3 shows that the two tumors that did not disappear were included in the groups of 750 and 1000 mW/cm^2^, and both cases were irradiated for 111 s.

**Fig. 3.**
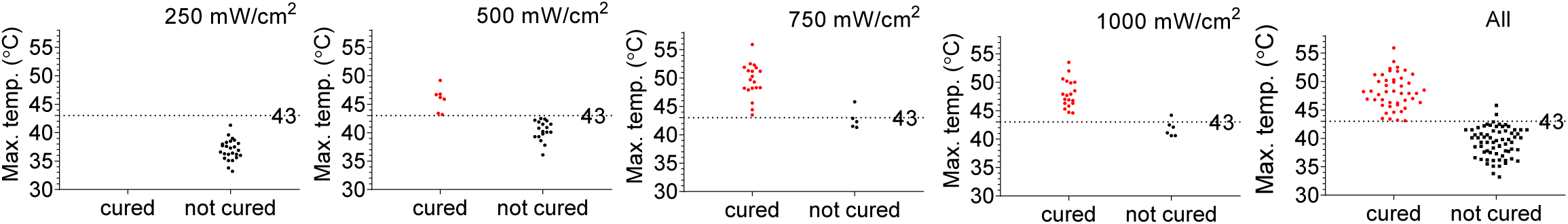
Scatter plots of the maximum temperature in individual tumor during NIR irradiation. Each plot is divided into two groups (“cured” and “not cured”). Fluence rate (mW/cm^2^) is shown in each graph. The results indicate that tumors are cured at a temperature above 43°C regardless of the fluence rate.

The results suggested that tumor cure depends on the tumor temperature during NIR irradiation. We have developed a thermal sensor circuit-based NIR laser irradiation system using a non-contact thermometer in order to keep the temperature of the irradiated target constant during irradiation (Nomura et al., 2017). We carried out a NIR irradiation experiment using the temperature-feedback laser system. Using this system, tumors were irradiated at 40, 41, 42 or 43°C for 333 s, and tumor size was measured up to 21 days after irradiation. The results showed that the tumor was not cured at a temperature below 41°C. However, 3 of 5 tumors were cured at 42°C and all of the tumors were cured at 43°C (Table 1).

**Table 1.**
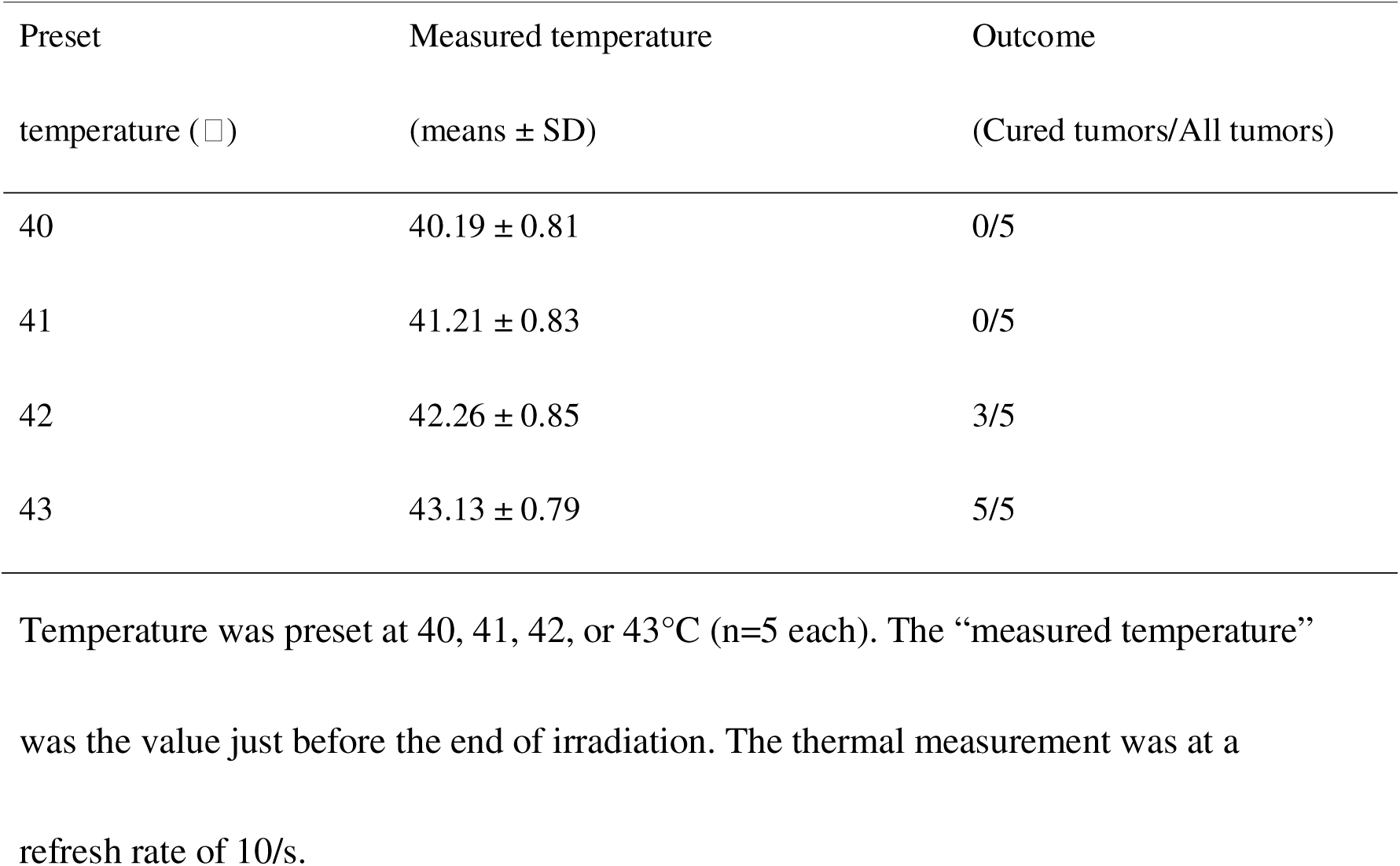
Outcomes of treatment of tumors with the temperature-controlled NIR laser irradiation system in intradermal tumor model mice administered ICG lactosome (8.8 mg/kg) at 48 h before the laser irradiation.

### Negligible increase in temperature at the irradiated non-tumor site

To confirm that ICG lactosome does not accumulate in tissue other than the tumor tissue and that the increase in temperature at non-tumor sites is negligible, temperatures of the tumor site and non-tumor site were measured during NIR laser irradiation (Fig. 4A). The increases in temperature (means ± SD) at the tumor site from the initial temperature to maximum temperature at 250, 500, 750 and 1000 mW/cm^2^ were 5.6 ±1.0, 10.6 ± 1.2, 13.6 ± 3.1, 19.0 ± 1.9°C, respectively, while those at the non-tumor site were 0.2 ± 0.1, 1.6 ± 0.2, 2.7 ± 0.4, 4.6 ± 0.3°C, respectively (Fig. 4B): the higher the fluence rate was, the larger was the difference in temperature increase. We also compared the differences in temperature increases at non-tumor sites with and without injection of ICG lactosome. In the case of ICG lactosome injection, the non-tumor site was irradiated at 48 hours after the injection. A slight increase in temperature during NIR irradiation was observed, and the increase in temperature was correlated with increase in fluence rate (Fig. 4C, red circles). However, the increase in temperature at the non tumor site in the case of ICG lactosome injection was not significantly different from that at the non-tumor site in the case of no ICG lactosome injection (Fig. 4C, black circles). These results suggest that there is little accumulation of ICG lactosome at sites other than the tumor site.

**Fig. 4.**
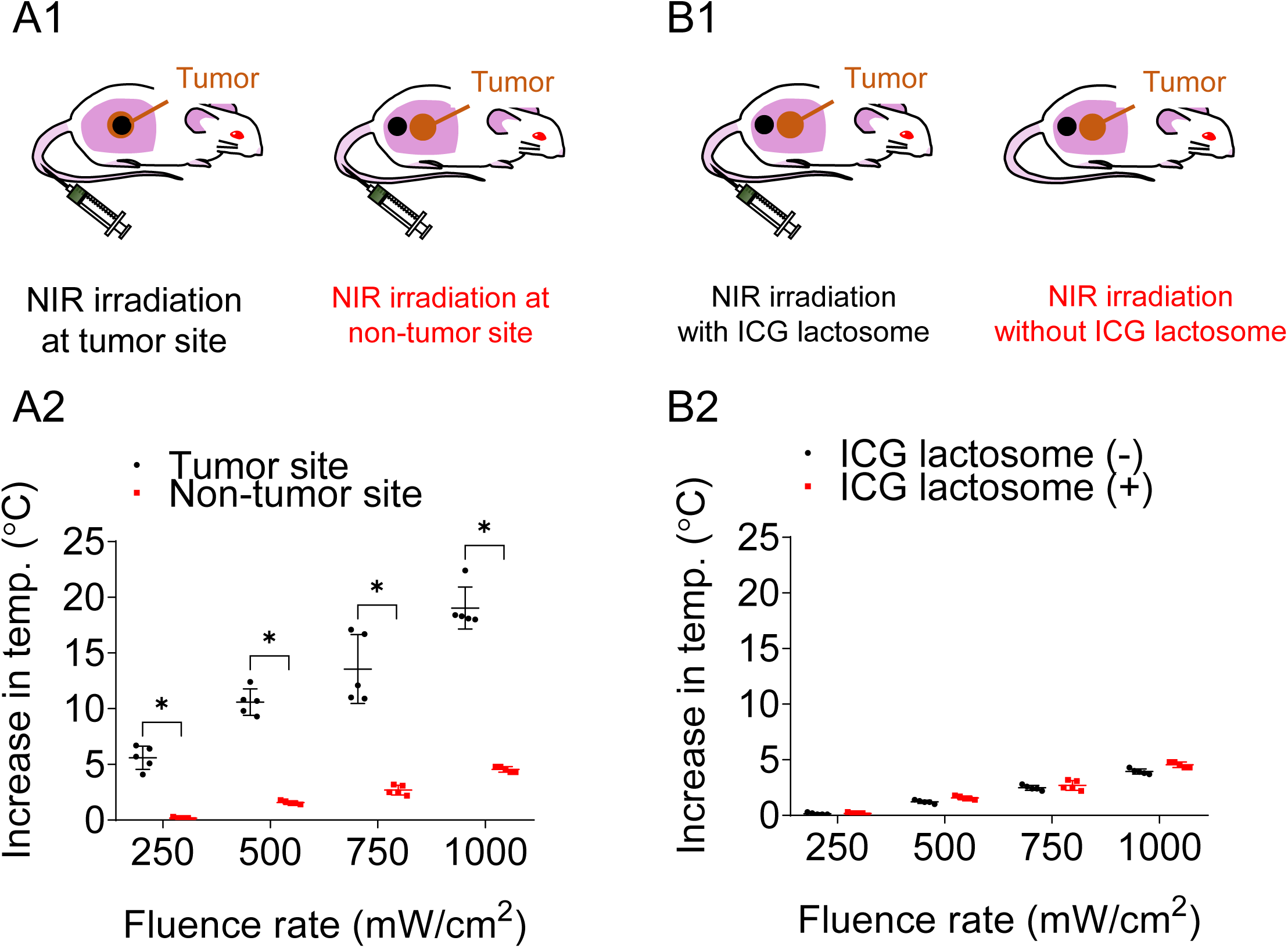
Increase in temperature at the spot of NIR irradiation on the tumor site or at that on a non-tumor site in mice with ICG lactosome administration (A1, A2). A1. Schema of the NIR irradiation point (black circle); A2. Scatter plots of increase in temperature at the tumor site and non-tumor site with ICG lactosome administration. With ICG lactosome administration, the higher the fluence rate was, the greater was the difference in increases in temperature at the tumor site and non-tumor site. (*P < 0.01, n= 5 each) Increase in temperature at the spot of NIR irradiation on a non-tumor site in mice with or without ICG lactosome administration (B1, B2). B1. Schema of the NIR irradiation point (black circle); B2. Scatter plots of increase in temperature at a non-tumor site with and without ICG lactosome administration. Increase in temperature at the non-tumor site with ICG lactosome administration is hardly distinguishable from that without ICG lactosome administration.

## DISCUSSION

This study showed that ICG lactosome selectively accumulated in the tumor, and it was also found that the photothermal reaction generated by the combination of ICG lactosome and NIR light suppressed the tumor growth. However, it was found that fluence rate and time of laser irradiation were not determinants for tumor cure. It was shown that tumors disappeared when the maximum temperature reached or exceeded 43°C during NIR irradiation. In addition, the increase in temperature due to NIR irradiation was negligible at non-tumor sites even when ICG lactosome had been injected. Thus, ICG lactosome facilitates a photothermal effect only at the tumor site without increasing the temperature around the tumor (non-tumor sites).

The results suggest that thermal dosimetry during NIR irradiation is a more useful index than fluence rate or irradiation time for prediction of tumor cure. We verified this concept by combining ICG lactosome and a temperature-controlled NIR laser system that can alter the output power of light to keep the temperature of the irradiated tumor constant. A temperature of 43°C was revealed to be critical for cure of an irradiated tumor.

Most of the studies on PTT have aimed to improve the characteristics of photo-absorbing agents such as strong absorbance, high photothermal conversion efficiency, and good photostability (Ali et al., 2017, Chen and Cai, 2015, Shanmugam et al., 2014, Song et al., 2015). Although this direction may be important for PTT, determination of the appropriate threshold temperature at which tumors are reliably cured might be more important. The distribution of nanomaterials in a tumor varies due to the heterogeneity of cancer tissue (Kronig et al., 2015), the abnormal structure of tumor blood vessels (Jackson et al., 2007) and the tumor interstitial pressure (Wu et al., 2014). Due to the heterogeneity of the tumor, it is natural to expect the outcomes to be erratic in the case of the conventional laser therapy in which the irradiation parameters (fluence rate and irradiation time) are set before laser irradiation. PTT with thermal dosimetry during laser irradiation can overcome the problems due to tissue heterogeneity and has the advantage of achieving highly reliable therapeutic effects.

Some reports indicating cure of tumors when the temperature was raised to 43°C or higher support the above-mentioned concept. Yoo et al. reported that hyperthermia induced by the application of an alternating magnetic field was able to cure a tumor at a temperature above 43°C (Yoo et al., 2013). O’Neal et al. reported that PTT with gold-silica nanoshells cured tumors by raising the temperature of the tumor up to 50°C (O’Neal et al., 2004).

Thermal dosimetry-assisted PTT may be useful for clinical use not only to effectively cure tumors but also to not damage normal tissues. It would be convenient for clinicians to predict the therapeutic effect during irradiation. Generally, the protocols of laser therapies including photodynamic therapy are set before irradiation. Thus, even if individual tumor phenotypes are different, the tumor must be irradiated only with the fixed parameters (fluence rate and irradiation time) in the clinical setting.

There are some limitations in this study. First, the tumor model was an intradermal tumor model for monitoring surface temperature of the tumor by using a non-contact thermometer. The surface temperature has been shown to be correlated with the internal temperature in an intradermal tumor model (Zhu et al., 2016), although that needs to be examined in detail in the future. Second, colon cancer was examined in this study. However, in different types of tumor, the tissue structure, heat tolerance and therapeutic threshold temperature might be different from those in other cancer types.

## Conclusions

In conclusion, ICG lactosome was highly accumulated in the intradermal tumor model and when the surface temperature of the tumor exceeded 43°C, the tumor disappeared in this model regardless of the fluence rate or irradiation time. PTT using ICG lactosome with thermal dosimetry will be a novel and useful theranostic tool for cancer strategy and provide clinicians with more reliable cancer treatment.

## Materials and Methods

### Tumor-targeted photo-absorbent

The ICG lactosome is a micelle-based agent that was synthesized as previously reported (Makino et al., 2007, Tsujimoto et al., 2015a, Tsujimoto et al., 2014a). In brief, the ICG lactosome used in this study contained 22% of ICG– poly (L-lactic acid) and 78% of poly (sarcosine)–poly (L-lactic acid). Regarding ICG– poly (L-lactic acid) synthesis, the terminal end of poly (L-lactic acid) was chemically modified using ICG as follows: 1.0 mg ICG–OSu was added to a dimethylformamide (DMF) solution of the free amino group bearing poly (L-lactic acid) (2.46 mg), which contains an amino group designed as an indicator of sarcosine N-carboxyanhydride (NCA) polymerization during the synthesis of amphiphilic poly (sarcosine)–poly (L-lactic acid)-block copolymers. The reaction mixture was stirred at room temperature overnight under a light-shielding condition. The reaction mixture was purified by size-exclusion chromatography using a Sephadex LH–20 column (GE Healthcare Japan Corporation, Tokyo, Japan) with DMF as the eluent.

Chloroform solutions of PS–PLLA and ICG–PLAA were mixed at a ratio of 0.78:0.22. The solvent was removed under reduced pressure and the thin film that had formed was dissolved in 10 mM Tris–HCl buffer (pH 7.4). The resulting aqueous solution was purified by Sephacryl S-100 size-exclusion chromatography by elution with 10 mM Tris–HCl buffer (pH 7.4) to obtain ICG lactosome. Dynamic light scattering analysis revealed that the hydrodynamic diameters of ICG lactosome particles were 30–40 nm, and the zeta potential of ICG lactosome was −0.51 mV. Since poly (sarcosine)–poly (L-lactic acid) is comprised of a biodegradable polypeptide, its toxicity is negligible, thus suggesting its safe use in humans.

### Animals

Female Balb/c mice at 6 weeks of age (Japan SLC, Hamamatsu, Japan) were fed under specific pathogen-free conditions. All animal procedures followed the guidelines approved by the National Defense Medical College Animal Care and Use Committee.

### Cell lines

A murine Colon26 cell line (NLC26) stably expressing Nano-lantern (Saito et al., 2012) was established as previously described (Maday et al., 2008). The cells were cultured in DMEM medium (Sigma-Aldrich, St. Louis, MO) supplemented with 10 % heat-inactivated fetal bovine serum (Life Technologies, Carlsbad, CA), 100 U/mL of penicillin, 100 µg/mL of streptomycin, and 0.25 µg/mL of amphotericin B (Antibiotic-Antimycotic, Life Technologies) at 37°C in 5% CO2 with 95% humidity.

### Establishment of a mouse intradermal tumor model

To establish the experimental intradermal tumor model, mice were injected with 0.5×10^6^ NLC26 cells suspended in 50 µL of phosphate-buffered saline into the skin of the right back under anesthesia. A combined anesthetic, prepared with 0.3 mg/kg of medetomidine 4.0 mg/kg of midazolam, and 5.0 mg/kg of butorphanol, was used (Kawai et al., 2011).

### *In vivo* Imaging

For visualization of the tumor localization, 100 μL of coelenterazine h (Wako Pure Chemical Industries Ltd., Osaka, Japan) solution (2.5mg/mL) was administered intravenously through the retro-orbital sinus of an anesthetized mouse. A luminescence image was taken using the IVIS, followed by acquisition of a white-and-black image and overlay of the pseudo-color image representing the spatial distribution of photon counts produced by Nano-lantern within the tumor.

For visualization of fluorescence from ICG lactosome accumulated in the tumor, 8.8 mg/kg ICG lactosome corresponding to 281 µM in ICG concentration was injected intravenously through the retro-orbital sinus of an anesthetized mouse 48 h before imaging. The fluorescence image was taken using the IVIS: the tumor was illuminated with a 780-nm excitation light and the fluorescence was captured using an 845-nm band-pass filter with an irradiation time of 500 ms. Fluorescence imaging was performed before and after NIR laser irradiation.

### NIR laser irradiation and thermal dosimetry

ICG lactosome was intravenously administered 4 or 5 days after intradermal inoculation of the cancer cells. At that time, the tumor of the right back was evident because of the cancer cell growth, and the size was approximately 200 mm^3^. Forty-eight hours after administration of ICG lactosome, the tumor was irradiated using a fiber-coupled laser system with a laser diode at 808 nm (model FC-W-808, maximum output: 10 W; Changchun New Industries Optoelectronics Technology Co., Ltd., Jilin, China). The fiber probe was placed just above the tumor so that the irradiated light spot was 1.0 cm in diameter, corresponding to a spot area of 0.79 cm^2^. The fluence rate was set at 250, 500, 750 or 1000 mW/cm^2^ and the irradiation time was set 111, 222, 333, 666 for 1000 s. The temperature of the tumor surface was measured with an infrared radiation thermometer. Tumor size was repeatedly measured using a digital caliper until the 21st day after irradiation, and tumor volume was calculated by the following equation: (longitudinal size) × (transverse size) × (transverse size) ×4/3π. Tumor disappearance was judged using histochemical images and luminescent images obtained by using the IVIS. Tumor cure was defined as the case disappearance of luminescence from coelenterazine h and no observation of a tumor histologically.

### Temperature controlled NIR laser irradiation experiments

Since the results obtained by PTT suggested that the peak of temperature of the tumor during NIR irradiation is the most reliable index to predict tumor shrinkage, we conducted animal experiments using a system that can control the temperature of the tumor. In order to keep the temperature constant, we used a NIR laser system for which the irradiation power is modulated by a temperature feedback control circuit that we previously reported (Nomura et al., 2017). The temperature was set 40, 41, 42 or 43°C (n = 5 in each). The irradiation time was set at 333 s.

### Statistical methods

Data are presented as the means ± standard deviation (SD) in Fig. 2B and Table 1. Statistical analyses were performed using the two-sample Kolmogorov-Smirnov test or chi square test with Fisher’s exact test where appropriate. Data are presented as the means ± standard error (SE) in Fig. 4.. Statistical analyses were performed using the two-stage linear step-up procedure of Benjamini, Krieger and Yekutieli, with Q = 1%, where appropriate. Calculations were performed using GraphPad Prism8 software for Windows. A p value of <0.05 was considered statistically significant.

## Acknowledgments

The authors would like to thank W. Kayukawa, T. Takee and M. Ushida for technical assistance with the experiments. This work was supported by the Japan Society for the Promotion of Science (Kakenhi) Grant Number 17H02114, the Japanese Foundation for Research and Promotion of Endoscopy, and the Nakayama Cancer Research Institute.

## Author contributions

S.N., H.M., H.T., and Y.M. designed the research; S.N., M.H., I.H., E.O., A. S., E.T., and Y.M. performed the research; S.N., H.T., D.S., K.H., Y.K., H.U., and Y.M. analyzed data; and S.N., A.S., H.U., and Y.M. wrote the paper.

