## Supplemental Figure for "Highly reliable, targeted photothermal cancer therapy combined with thermal dosimetry using indocyanine green lactosome"

Corresponding author

**This PDF file includes:**

Figs. S1 to S2

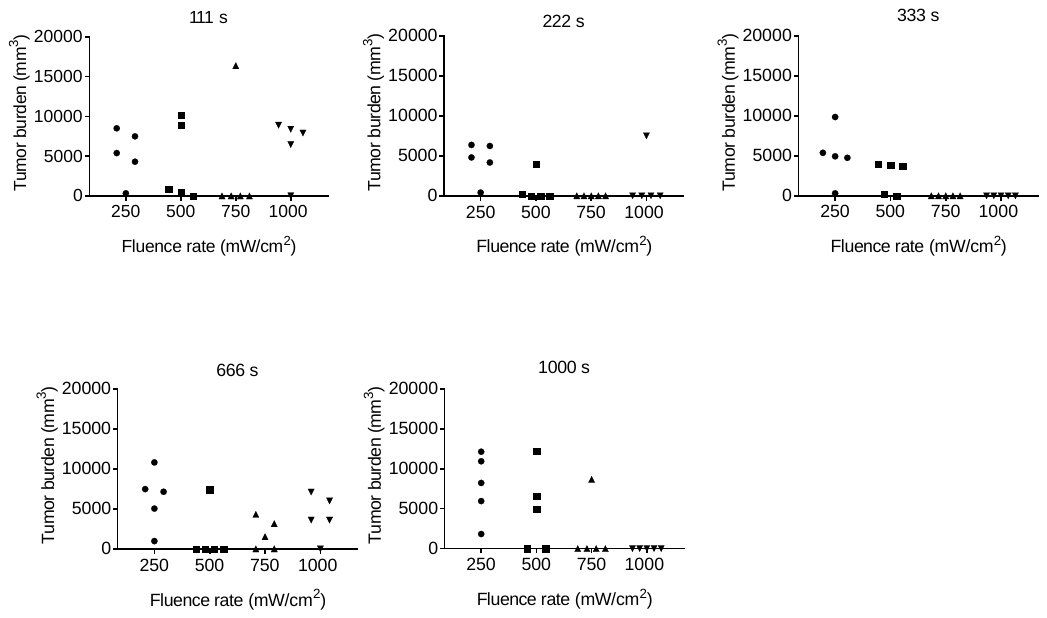

Fig. S1. Scatter plots of the tumor burdens in mice when irradiated at a fluence rate of 250, 500, 750 or 1000 mW/cm2. Data were obtained on the 21st day after irradiation (n=5 each).

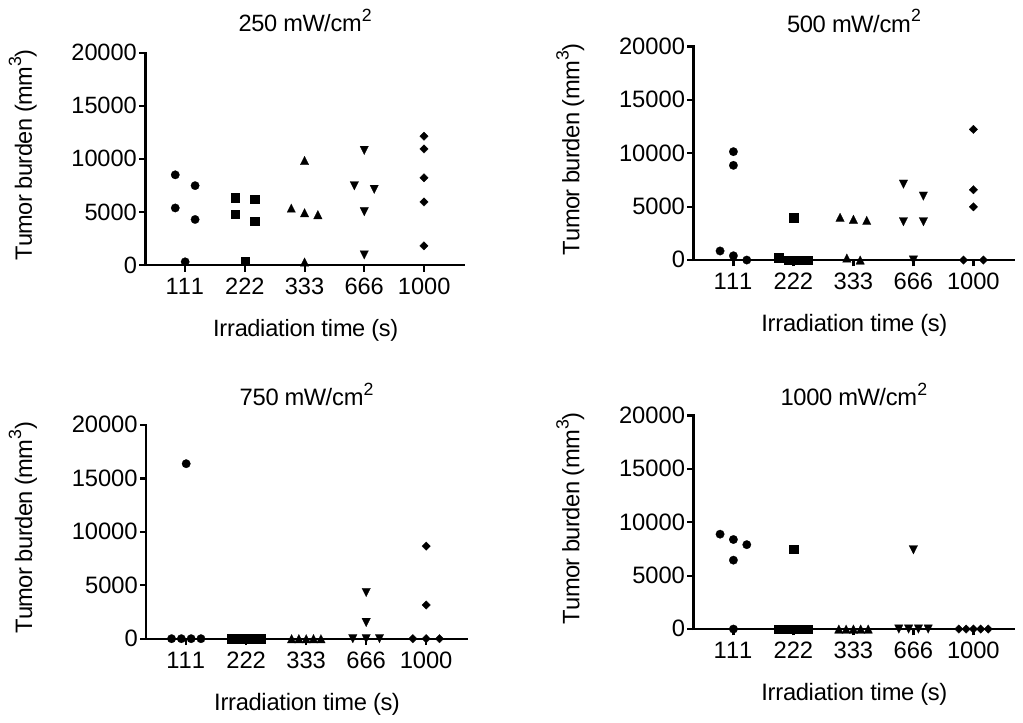

Fig. S2. Scatter plots of the tumor burdens in mice when irradiated with irradiation time of 111, 222, 333, 666 or 1000 s. Data were obtained on the 21st day after irradiation (n=5 each).
